# Phenotypic plasticity during diel cycling hypoxia in Arctic char (*Salvelinus alpinus*)

**DOI:** 10.1101/2022.12.24.521867

**Authors:** Loïck Ducros, Mohamed Touaibia, Nicolas Pichaud, Simon G. Lamarre

## Abstract

Oxygen concentration naturally fluctuates in aquatic environments. Due to increased eutrophication caused by anthropic activities, this phenomenon could be amplified and result in a daily cycle of alternating normoxic and hypoxic conditions. At the metabolic level, lack of oxygen and reoxygenation can both have serious repercussions on fish due to fluctuations in ATP supply and demand and an elevated risk of oxidative burst. Thus, fish must adjust their phenotype to survive and equilibrate their energetic budget. However, their energy allocation strategy could imply a reduction in growth which could be deleterious for their fitness. Although the impact of cyclic hypoxia is a major issue for ecosystems and fisheries worldwide, our knowledge remains however limited. Our objective was to characterise the effects of cyclic hypoxia on growth and metabolism in fish. We monitored growth parameters (specific growth rate, condition factor), hepatosomatic and visceral indexes, relative heart mass and hematocrit of Arctic char (*Salvelinus alpinus*) exposed to thirty days of cyclic hypoxia. We also measured the hepatic protein synthesis rate, hepatic triglycerides as well as muscle glucose, glycogen and lactate, and quantified hepatic metabolites during this treatment. Arctic char appeared to acclimate well to oxygen fluctuations. The first days of cyclic hypoxia induced a profound metabolome reorganisation in the liver. However, fish rebalanced their metabolic activities and successfully maintained their growth and energetic reserves after one month of cyclic hypoxia. These results demonstrate the impressive ability of fish to cope with their changing environment.

**Summary statement:** This study characterizes the metabolic adjustments performed by Arctic char when coping with one month of cyclic hypoxia. Fish reached a new phenotype by defending their growth and energy stores.

## Introduction

The intensification of anthropic activities is currently amplifying eutrophication in aquatic environments (Jickells, 2005; Le Moal *et al*., 2019; Stets *et al*., 2015) resulting in an increase in algae biomass (Jickells, 2005; Le Moal *et al*., 2019; Stets *et al*., 2015). As a result of photosynthesis and respiration, the concentration of dissolved oxygen increases during the day and drops drastically at night, creating a daily cycle of alternating normoxic and hypoxic conditions (Miyamoto *et al*., 2019; Odum, 1956). Both the lack of oxygen and the reoxygenation may impose a challenge on aquatic organisms such as fish.

Decreased oxygen concentration in the aquatic environment can lead to a reduction of systemic PO_2_ and hypoxemia (Farrell and Richards, 2009). One of the first responses is hyperventilation resulting from increased ventilatory amplitude or frequency (Eom and Wood, 2020; Perry *et al*., 2009). Fish may also improve oxygen transport by increasing hematocrit (Jia *et al*., 2021) or via the Root effect (Rummer and Brauner, 2015). If oxygen supply is insufficient, tissues become hypoxic (Samuel and Franklin, 2008) which leads to a decrease of ATP production through oxidative phosphorylation. To avoid a mismatch between ATP supply and demand, anaerobic glycolysis is activated (Abdel-Tawwab *et al*., 2019; Speers-Roesch *et al*., 2013). Fish may also switch into an hypometabolic state and reorganize their metabolism (Nilsson and Renshaw, 2004; Richards, 2009). Multiple metabolic pathways are inevitably affected. Besides glycolysis and TCA cycle, amino acids, purine and lipid metabolism are also impacted in a tissue-specific manner (Dahl *et al*., 2021; Farhat *et al*., 2021). The liver is involved in carbohydrate, lipid, and protein metabolism (Charlton, 1996; Rui, 2014) and as such, plays a central role in energy metabolism. During hypoxia, carbohydrates and lipids are mobilized to produce ATP while the rate of protein synthesis is depressed to reduce energy demand (Cassidy and Lamarre, 2019; Cassidy *et al*., 2018; Lardon *et al*., 2013; Li *et al*., 2018). Moreover, hypoxia may decrease glycogen and lipid stores (Cadiz *et al*., 2018; Gracey *et al*., 2011). Glucogenic amino acids could also be catabolized to provide substrates for gluconeogenesis (Ding *et al*., 2020). Due to this high pressure on metabolism, hypoxia may cause an impairment in reproduction, growth and survival of fish (Aksakal and Ekinci, 2021; Landry *et al*., 2007; Pichavant *et al*., 2001).

The return to normoxia following a hypoxic bout could be seen as a welcomed event for fish. However, it is associated with effects similar to ischemia-reperfusion (Li and Jackson, 2002). The return of oxygen causes an oxidative burst (Loor *et al*., 2011) due to the production of reactive oxygen species (ROS) by mitochondria (Li and Jackson, 2002). Chouchani *et al*. (2016) demonstrated a mechanism that explains this oxidative burst. During the hypoxic phase, the lack of oxygen as the final electron acceptor results in a reduction of the electron transport system (ETS), especially at the level of coenzyme Q. Fumarate then captures the excess electrons and is reduced to succinate which serves as electron storage (Chouchani *et al*., 2016). During reoxygenation, the massive supply of electrons by oxidation of the accumulated succinate causes a reverse transport of electrons to complex I, resulting in a significant production of ROS and associated damages to lipids, proteins and DNA (Kehrer, 1993).

Although hypoxia is becoming a major issue for ecosystems and fisheries worldwide, our knowledge is limited regarding the impact of cyclic hypoxia. In this study, we explored the effects of multiple hypoxia-reoxygenation cycles on the growth, physiological conditions, and metabolism of Arctic char. We hypothesized that fish would have to alter their phenotype and use their energetic reserves to overcome exposure to cyclic hypoxia. We predicted that Arctic char experiencing chronic cyclic hypoxia would have a reduced growth rate and a lower condition factor compared to control fish, as well as reduced energetic reserves. Moreover, we predicted that to compensate for the slower rate of protein synthesis during the hypoxic periods, a compensation of that rate would be observed following reoxygenation. Finally, we predicted that these metabolic adjustments would be reflected in the fish metabolome. To test this hypothesis, we exposed Arctic char to a month of cyclic hypoxia and measured growth parameters, protein synthesis rate and triglycerides (TAG) in the liver, carbohydrate reserves in white muscle, and characterized the liver metabolome.

## Materials and methods

### Animals

Arctic char were obtained from Valorēs Inc. (Shippagan, NB, Canada). The fish were kept in 250 L tanks with air saturated dechlorinated water supplied at approximately 16°C, in a 14:10 light cycle (220 lux). The fish were fed *ad libitum* three times a week with a commercial diet (Nutra RC 80A, Skretting, Bayside, NB, Canada). All animal experiments were approved by the Institutional Animal Care Committee (UdeM 19-12). At least two weeks before the experiment, the fish were individually tagged with 8 mm x 1.4 mm FDX PIT tags (Oregon RFID, Portland, USA).

### Experimental design

For the cyclic hypoxia experiments, we used a recirculation system composed of two independent units (Multi-stressor system, Aquabiotech Inc., QC, Canada). A total of 240 fish (16.56 g ±0.21, 12.90 cm ±0.06) were randomly transferred into ten 9 L tanks (24 fish per tank). The fish were fed 0.5% biomass each day and fasted 24 hours prior sampling. After fourteen days of acclimation in air-saturated water at 16°C, half of the tanks were exposed to cyclic hypoxia for 30 days. The oxygen concentration was reduced to 20 % air saturation between 8:00 pm and 6:00 am (hypoxic phase) and maintained at 100% air saturation for the rest of the day (normoxic phase). In each tank, four fish were sampled at 9:00 am on days 1, 5, 10 and 30 (D1, D5, D10, D30). The tank was considered as the experimental unit (N =5 per condition). The density of fish in each tank was kept constant using a sliding plexiglass wall to limit the formation dominance hierarchy. The fish were weighted and measured after being euthanized by a blow to the head. A blood sample was taken and centrifuged at 1,000×*g* for 5 minutes at 4 ° C in a hematocrit capillary. Liver, heart, digestive tract, and white muscle were quickly dissected, frozen in liquid nitrogen, and stored at -80°C until further experiments. The hepatosomatic index, visceral index and relative heart mass were determined according to the following equations:

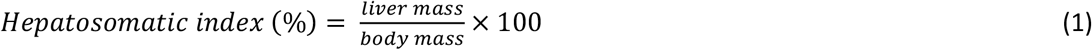

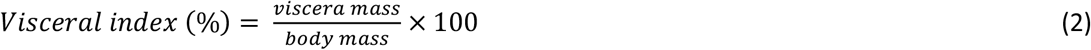

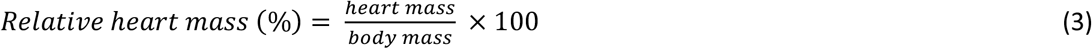

On day 0, 10, 20 and 30 (D0, D10, D20, D30), all fish were anesthetized in a 50 mg.mL-1 benzocaine solution, weighted, and measured. The specific growth rate (SGR) in length and mass as well as the condition factor (CF) were calculated according to the following equations:

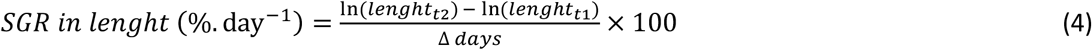

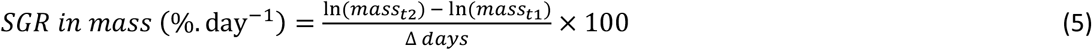

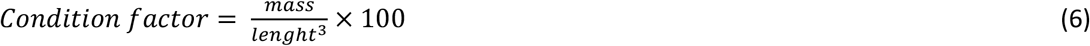

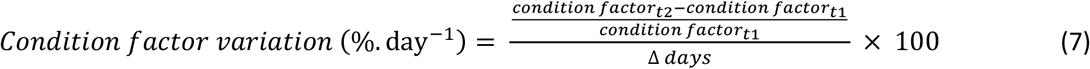

### Energy stores

White muscle glucose, glycogen, and lactate

Approximately 40 mg of frozen white muscle (2 fish processed per tank per condition) were homogenized by ultrasonication (Q55 Sonicator, Qsonica Inc., USA) in 5 volumes of 1 M perchloric acid (PCA; A228 Fisher Scientific, USA). The samples were neutralized by adding KHCO_3_. Glucose was assayed in a 250 mM imidazole buffer supplemented with 5 mM MgSO_4_·7H_2_O, 10 mM ATP, 1 mM NADP+, pH 7.8. Following the addition of G6PDH and hexokinase in excess, absorbance at 340 nm was recorded. Glucose concentration was determined from a standard curve and reported in (μmol.g-1 tissue). For glycogen, the neutralized samples were incubated 2 hours at 37 °C with amyloglucosidase (50 U.mL-1) in acetate buffer (0.4 M, pH 4.8). Glucose concentration was then determined as above, and glycogen concentration was reported in μmol glycosyl units per g tissue. Lactate concentration was determined as described in Brandt *et al*. (Brandt *et al*., 1980). Briefly, samples or standards were added to 10 volumes of assay buffer (320 mM glycine, 320 mM hydrazine, 2.4 mM NAD+, pH 9.5). Lactate concentration was determined by reading absorbance at 340 nm after a 150-minute incubation at 25°C with LDH in excess. Results are presented as means ± se and expressed as μmol.g-1 tissue.

### Liver triglycerides

Lipids were extracted from approximately 20 mg of frozen liver (2 fish processed per tank per condition) using chloroform/methanol extraction (Bligh and Dyer, 1959). Extracted lipids were resuspended in a 95% ethanol. Lipid concentration was determined using a commercial kit (Infinity Reagent, Thermo Scientific, USA) and reported as means ± se and expressed as μmol glycerol units per g tissue.

### Protein synthesis

The fractional rate of protein synthesis (*Ks*) was determined following the flooding dose method developed by Garlick *et al*. (Garlick *et al*., 1980) and modified according to Lamarre *et al*. (2015). Three fish per tank per condition were injected with a 150 mM phenylalanine (PHE) solution containing 50% ring-[D_5_]-L-phenylalanine ([D_5_]-PHE, 98%, Cambridge Isotope Laboratories, Inc. Andover, USA) at a dose of 1 mL per 100 g of body mass and returned to their tanks for an incorporation period of three hours. The fish were then sacrificed, and the liver was flash frozen and kept at -80°C. Approximately 20 mg of frozen tissue were homogenized in 0.2 M PCA by ultrasonication and centrifuged. The supernatant, which contains protein-free amino acid pool (FP), was collected and stored at -20 °C. The pellet containing the protein pool (PP) was washed three times in 0.2 M PCA. The pellet was finally washed with 1 mL of acetone to remove excess lipids before being hydrolyzed in 6 M HCl at 110 °C for 18 hours. PP and FP phenylalanine was extracted using solid phase extraction as described in Lamarre *et al*. and Cassidy *et al*. (Cassidy *et al*., 2016; Lamarre *et al*., 2015). The extracted amino acids were derivatized using pentafluorobenzyl bromide (PFBBr). Briefly, 50 μL of sample were incubated 45 minutes at 60 °C with 20 μL of phosphate buffer (0.5 M, pH 8.0) and 130 μL of PFBBr (100 mM in acetone). Derivatized amino acids were extracted in 330 μL of hexane, transferred to low volume inserts before analysis. GC-MS analyses were performed using an Agilent gas chromatograph (model 6890N) interfaced to a single quadrupole inert mass selective detector (MSD, model 5973) as previously described by Lamarre *et al*. (2015). *Ks* (%.day-1) was calculated according to the Eq. 8:

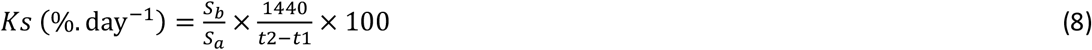

Where *S*_*a*_ is the FP enrichment (*S*_*a*_ = [D_5_]-PHE / (PHE + [D_5_]-PHE)), *S*_*b*_ the PP enrichment (*S*_*b*_ = [D_5_]-PHE / (PHE + [D_5_]-PHE)) and *t* the incorporation time in min.

### Metabolite extraction and quantification

Approximately 50 mg of frozen liver (of fish that were not used in the protein synthesis assay) were homogenized in 250 μL of cold acetonitrile with an ultrasound homogenizer on ice. 250 μL of cold water were then added and the solution was homogenized by sonication on ice. After a 5 min 10,000×*g* centrifugation at 4°C, 400 μL of the supernatant containing hydrophilic metabolites were transferred to a borosilicate tube and evaporated 1 hour under a nitrogen stream. The samples were stored at -80 °C until further analysis.

Just before 1H NMR analysis, samples were resuspended in 900 μL of D_2_O and 100 μL of 4,4-dimethyl-4-silapentane-1-sulfonic acid (DSS, internal calibrator, 0.5 mM final concentration). Approximately 700 μL were transferred to a 5 mm NMR tube. The 1H-NMR spectra were recorded on a Bruker Advance III 400 MHz spectrometer at 298 K. Analyses were performed with the *noesypr1d* pulse sequence. One-dimensional spectra were obtained after 128 scans of 64,000 data points. The spectral width was set at 12 KHz, the acquisition time at 6.6 seconds, and the recycle delay at 1 second per scan. Correction and calibration of the spectra were performed using the Chenomx NMR Processor (Chenomx Inc., Edmonton, AB, Canada) and analyzed using the Chenomx NMR Profiler (Chenomx Inc., Edmonton, AB, Canada) and the Human Metabolome Database (HMDB) (Wishart *et al*., 2018).

### Statistics

Statistical analyses were performed with the R software (R Core Team, version 4.1.0, 2021). Data were fitted to a mixed linear model with treatments (normoxia, cyclic hypoxia) and sampling day (or periods) as fixed factors and experimental tank as random factor. Normality and homoscedasticity of residuals were checked with a Shapiro-Wilk test and a Levene test, respectively. When necessary, data transformation was applied. Two-way ANOVAs were performed to test the effects of treatment and sampling day (or periods) on growth, condition factor (CF) variation, energy store or Ks. When a significant interaction was detected, a *post hoc* test was performed using the least squares mean method with p values adjusted by the Tukey method. The significance threshold was set to p < 0.05. A two-sample t-test was performed to test the effect of treatment on SGR and CF variation after 30 days of exposure.

Statistical analyses of the metabolomics data were performed using MetaboAnalyst 5.0 (https://www.metaboanalyst.ca/). Metabolite concentrations were analyzed using a partial least squares discriminant analysis (PLS-DA) (Chong *et al*., 2019) to determine the variable importance to the projection (VIP) score. Data were log transformed and auto scaling (*i*.*e*. mean-centered and divided by the standard deviation of each variable) was applied in order to minimize possible differences in concentration between samples. Metabolites that were assigned a VIP score > 1.0 were considered important in the PLS-DA model (Chong *et al*., 2019) and were further analyzed for specific differences (two-sample t-test).

## Results

### Growth

We monitored several growth parameters of fish exposed to thirty days of normoxia or cyclic hypoxia. Average SGR in length and mass were not significantly different between the two groups at the end of the experiment (Table S1). The condition factor (CF) remained overall stable and did not differ between treatments (Table S1). Analyzing the results using periods of ten days revealed that SGR in length was stable (Fig. 1A), while SGR in mass changed significantly over time (F_2,16_=20.39, p < 0.001; Fig. 1B). There was no significant difference in the normoxic group, however the cyclic hypoxia group gained less mass during the first ten days (D0-D10) than during the D10-D20 (p < 0.001) and D20-D30 periods (p = 0.001).

**Figure 1.**
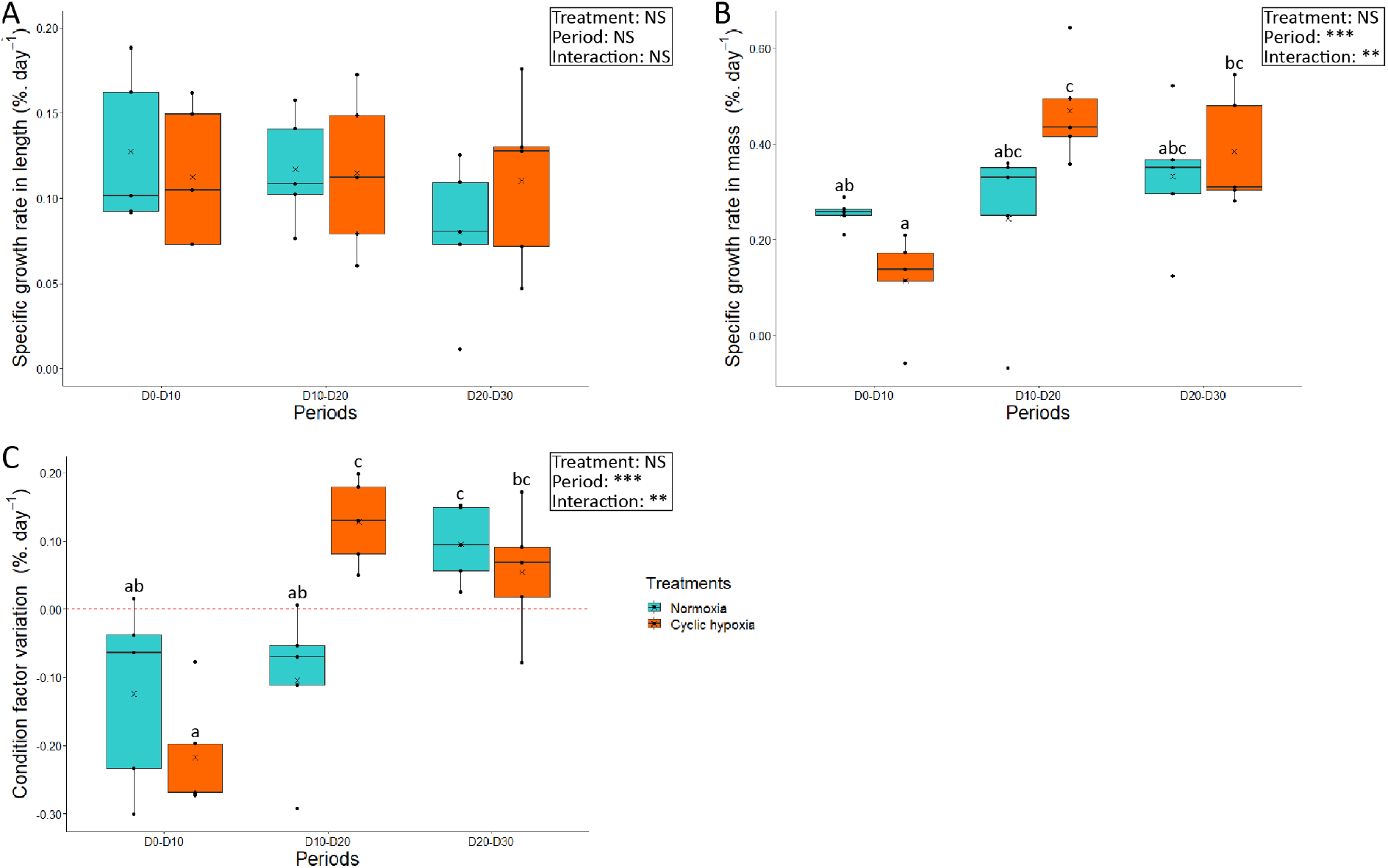
Specific growth rate in length (A) and mass (B) and condition factor variation (C) of Arctic char for each ten-day period, between day 0 and day 30, after fish were either maintained in normoxia or exposed to cyclic hypoxia. Box plots indicate the median (—), the mean (×), 25^th^ and 75^th^ percentiles (box), 95% range (|) and each observation (●). Two-way ANOVA results are shown as NS (non-significant), ** (p-value < 0.01), *** (p-value < 0.001). Different letters indicate significant differences (post-hoc test, p-value < 0.05).

The condition factor variation was significantly affected by the period (F_2,16_=19.09, p-value < 0.001; Fig. 1C) similarly to SGR in mass. Both normoxia and cyclic hypoxia fish had a negative CF variation between D0 and D10. CF then increased between D10 and D20 for fish exposed to cyclic hypoxia compared to the first 10 days (p-value < 0.001). However, for fish in normoxia, CF only improved between D20 and D30 (p-value < 0.05) comparatively to the first 10 days. Furthermore, CF variation was significantly higher during cyclic hypoxia between D10 and D20 (p-value < 0.05) compared to the normoxic group.

### Oxygen supply

Overall, exposure to cyclic hypoxia resulted in a slightly higher hematocrit compared to normoxia (F_1,8_=7.76, p-value < 0.05) but the number of hypoxia cycles experienced by the fish did not affect the hematocrit (Fig. S1). Moreover, relative heart mass remained unaffected by cyclic hypoxia.

### Energy stores

The energy stores were not affected by cyclic hypoxia. In white muscle, neither glucose nor glycogen concentrations differed between the control and fish exposed to cyclic hypoxia (Table 1). Moreover, lactate was also stable.

**Table 1.** Means ±*se* and two-way Anova F-values for energy stores and lactate concentration of Arctic char at days 1, 5, 10 and 30, after fish were either maintained in normoxia or exposed to cyclic hypoxia. Significant results are shown as * (p-value < 0.05), ** (p-value < 0.01). Different letters indicate significant differences (post-hoc test, p-value < 0.05).

|  | Normoxia |  |  |  | Cyclic hypoxia |  |  |  | Two-way Anova (F-values) |  |  |
| --- | --- | --- | --- | --- | --- | --- | --- | --- | --- | --- | --- |
|  | D1 | D5 | D10 | D30 | D1 | D5 | D10 | D30 | Treatment<br>Num df= 1<br>Den df= 8 | Sampling day<br>Num df= 3<br>Den df= 24 | Interaction<br>Num df= 3<br>Den df= 24 |
| **White muscle** |  |  |  |  |  |  |  |  |  |  |  |
| Glucose <sup>†</sup> ($\mu\text{mol.g}^{-1}$) | 3.55 ±0.56 | 3.50 ±0.18 | 3.65 ±0.30 | 4.23 ±0.65 | 3.48 ±0.24 | 3.44 ±0.33 | 3.92 ±0.39 | 4.25 ±0.52 | 0.08 | 1.28 | 0.10 |
| Glycogen ($\mu\text{mol.g}^{-1}$) | 14.58 ±1.92 | 14.14 ±0.73 | 11.56 ±1.28 | 15.40 ±1.76 | 11.89 ±0.89 | 11.37 ±1.23 | 15.29 ±1.63 | 18.00 ±2.07 | 0.03 | 3.31* | 3.00 |
| Lactate ($\mu\text{mol.g}^{-1}$) | 34.61 ±5.51 | 32.88 ±1.12 | 35.29 ±2.32 | 28.67 ±3.47 | 36.59 ±4.80 | 37.20 ±3.52 | 32.64 ±3.39 | 32.31 ±2.77 | 0.42 | 0.88 | 0.41 |
| **Liver** |  |  |  |  |  |  |  |  |  |  |  |
| Triglycerides ($\mu\text{mol.g}^{-1}$) | 4.24 ±0.42 | 4.67 ±0.27 | 4.27 ±0.74 | 4.20 ±0.76 | 3.87 ±0.30 | 4.25 ±0.39 | 5.51 ±0.58 | 2.94 ±0.41 | 0.33 | 2.40 | 2.08 |
| Hepatosomatic index (%) | 1.04 ±0.08 <sup>ab</sup> | 1.01 ±0.09 <sup>ab</sup> | 1.16 ±0.09 <sup>ab</sup> | 1.01 ±0.05 <sup>ab</sup> | 0.97 ±0.05 <sup>a</sup> | 1.22 ±0.08 <sup>b</sup> | 1.11 ±0.04 <sup>ab</sup> | 1.04 ±0.03 <sup>ab</sup> | 0.12 | 5.24** | 4.89** |
| **Viscera** |  |  |  |  |  |  |  |  |  |  |  |
| Visceral index (%) | 5.17 ±0.31 | 5.50 ±0.42 | 4.97 ±0.18 | 4.84 ±0.23 | 5.50 ±0.37 | 5.10 ±0.37 | 5.23 ±0.25 | 4.93 ±0.25 | 0.03 | 4.21* | 2.61 |

<sup>†</sup> data Box Cox transformation applied

No differences were also detected in liver triglyceride concentrations (Table 1). The hepatosomatic index was unchanged between groups for the duration of the experiment, except for a slight increase between D1 and D5 in the cyclic hypoxia group. The visceral index was also not affected by the treatment throughout the experiment (Table 1).

### Protein synthesis

The rate of protein synthesis of the liver was not affected by cyclic hypoxia (Fig. 2). Contrary to our prediction, no compensation of Ks during reoxygenation occurred in the liver at the beginning of the exposure to cyclic hypoxia. Liver Ks decreased slightly over time in both groups (F_3,24_ = 5.67, p-value < 0.01).

**Figure 2.**
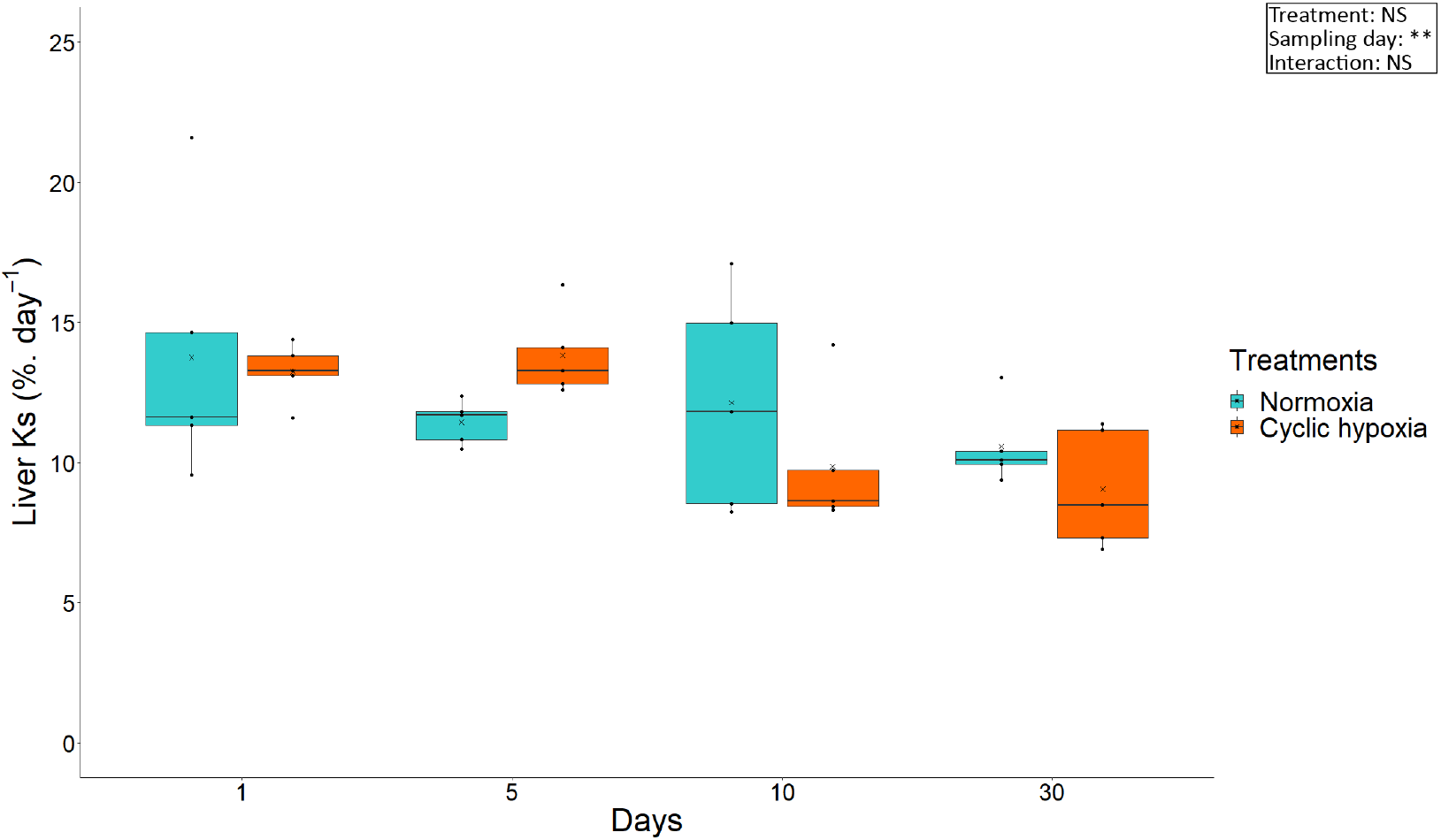
Liver Ks of Arctic char at days 1, 5, 10 and 30 after fish were either maintained in normoxia or exposed to cyclic hypoxia. Box plots indicate the median (—), the mean (×), 25^th^ and 75^th^ percentiles (box), 95% range (|) and each observation (●). Two-way ANOVA results are shown as NS (non-significative), ** (p-value < 0.01).

### Metabolomic analysis

A total of 68 metabolites were identified. Metabolites for which identification or quantification was not reliable, and metabolites from potential exogenous sources were removed. Finally, 49 metabolites were included in the analysis (Table S2). Metabolite concentrations were first analyzed over time in each treatment group (*i*.*e*., normoxia and cyclic hypoxia) using a PLS-DA. In normoxia, the two main components explained 10.6% and 7.6% of the total variation, respectively (Fig. S2A). This analysis revealed that D1 and D5 were more similar, while D10 differed slightly from the previous days. However, D30 was clearly separated from the early days. Twenty metabolites contributed to this metabolic change (Fig. S2B, VIP score > 1.0 for component 1). However, this model had low predictability (Q2) and accuracy (Table S3). During cyclic hypoxia, components 1 and 2 explained 17.7% and 9.4% of the variance, respectively (Fig. S3C). PLS-DA separated each sampling day with better segregation compared to the normoxic group. Twenty metabolites mainly drove this signature (Fig. S3D, VIP score > 1 for component 1) with moderate accuracy and predictability (Table S3). Considering that these changes over time largely resulted from the experimental conditions, we decided to focus on the differences between treatments at each sampling day.

Following one hypoxia-reoxygenation cycle, we observed a large separation between the normoxia and cyclic hypoxia groups on the first component, which represented 26.0 % of the total variation (Fig. 3A). This signature was driven by 18 metabolites (Fig. 3B). This model had high accuracy and predictability (Table S3). The main change was a 3.4-fold increase in alanine concentration in cyclic hypoxia (Fig. 3C). Furthermore, liver of fish exposed to cyclic hypoxia accumulated branched-chain amino acids (BCAAs; leucine, isoleucine, and valine) as well as beta-leucine, taurine, glucose, glyceraldehyde, and glucosamine-6-phosphate. Glutamate, carnitine, argininosuccinate and pyruvate levels were all significantly lower in the cyclic hypoxia group. Moreover, the concentrations of succinate and oxaloacetate were slightly lower but did not reach significance in the same group.

**Figure 3.**
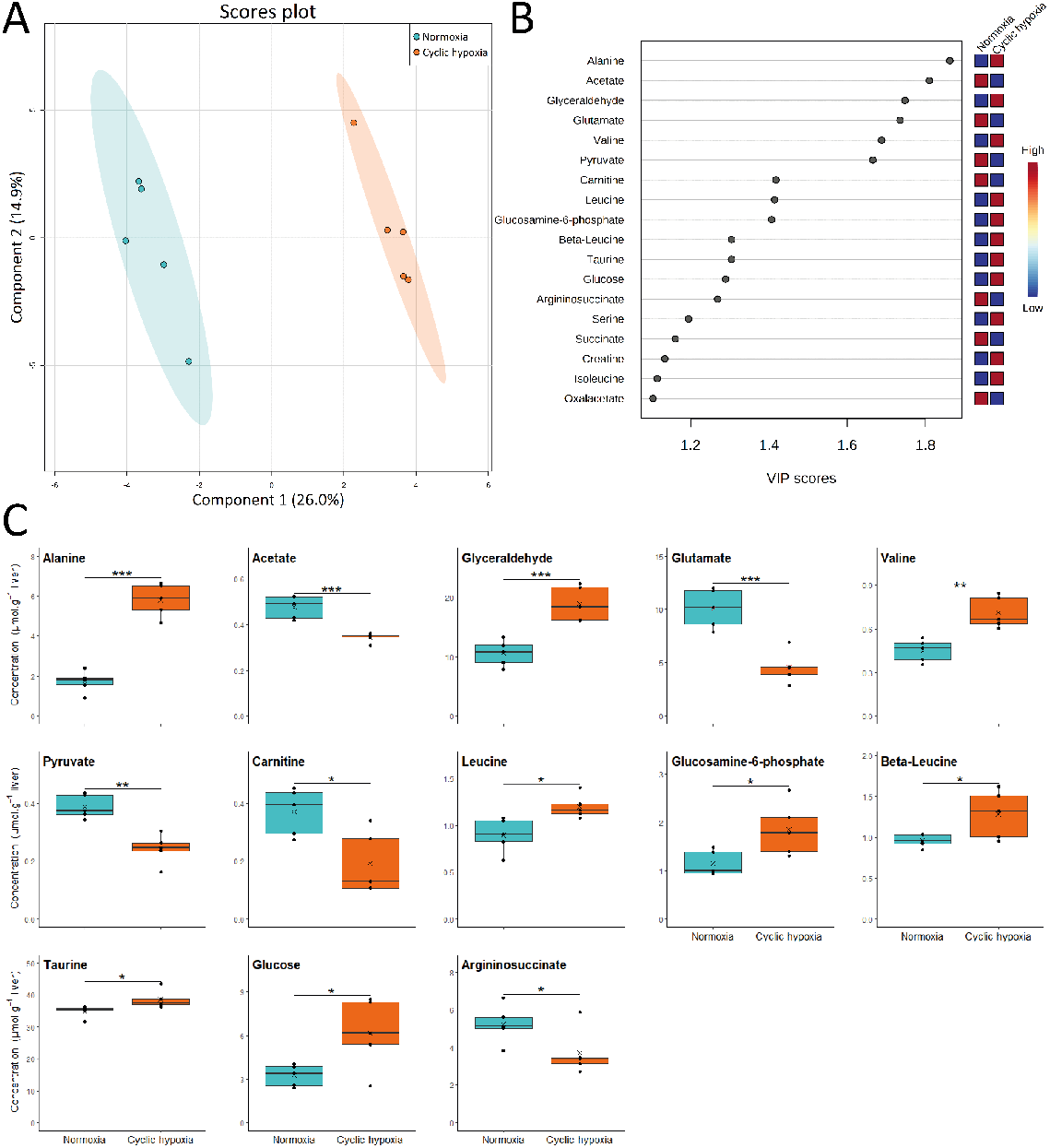
Metabolite profile in liver of Arctic char at day 1 after fish were either maintained in normoxia or exposed to cyclic hypoxia. A) PLS-DA 2D score plots of liver metabolites. Components 1 and 2 represent variance proportion and ellipses correspond to 95% confidence intervals for each treatment group. B) Variable importance in projection (VIP) scores of PLS-DA component 1 for liver metabolites which drive metabolic profile differentiation between the normoxic and cyclic hypoxic groups (VIP score > 1). Relative concentrations of corresponding metabolites are indicated by colored boxes on the right. C) Concentration of metabolites which differ significatively between normoxic and cyclic hypoxic fish. Box plots indicate the median (—), the mean (×), 25^th^ and 75^th^ percentiles (box), 95% range (|) and each observation (●). Two sample t-test results are shown as * (p-value < 0.05), ** (p-value < 0.01), *** (p-value < 0.001).

Five days of cyclic hypoxia resulted in a similar pattern. Differentiation of metabolic profile between treatments was also obvious with the first component explaining 24.5% of the variation (Fig. 4A). Eighteen metabolites led this pattern (Fig. 4B). The fish exposed to cyclic hypoxia had lower concentrations of glutamate and higher BCAAs than the normoxic fish (Fig. 4C). Cyclic hypoxia also led to a significant decrease in glycine concentration while the accumulation of glucose observed at D1 disappeared. Finally, the concentrations of malate, citrate, and fumarate were all elevated in the cyclic hypoxia fish.

**Figure 4.**
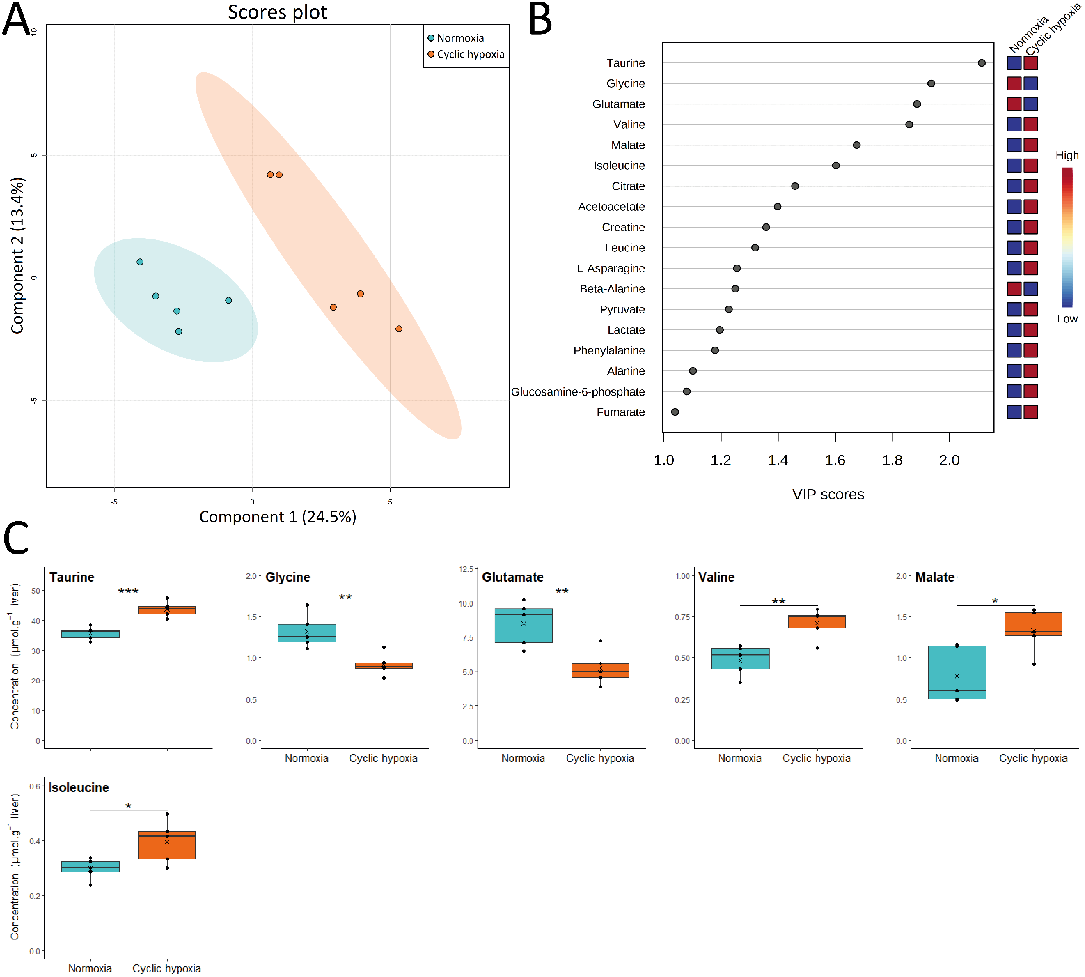
Metabolite profile in liver of Arctic char at day 5 after fish were either maintained in normoxia or exposed to cyclic hypoxia. A) PLS-DA 2D score plots of liver metabolites. Components 1 and 2 represent variance proportion and ellipses correspond to 95% confidence intervals for each treatment group. B) Variable importance in projection (VIP) scores of PLS-DA component 1 for liver metabolites which drive metabolic profile differentiation between the normoxic and cyclic hypoxic groups (VIP score > 1). Relative concentrations of corresponding metabolites are indicated by colored boxes on the right. C) Concentration of metabolites which differ significatively between normoxic and cyclic hypoxic fish. Box plots indicate the median (—), the mean (×), 25^th^ and 75^th^ percentiles (box), 95% range (|) and each observation (●). Two sample t-test results are shown as * (p-value < 0.05), ** (p-value < 0.01), *** (p-value < 0.001).

After ten days of exposure to cyclic hypoxia, the differences between normoxic and cyclic hypoxic fish were not as strong as the first days, as the variation explained by the first component (18.2%) was lower compared to D1 and D5 (26% and 24.5%, respectively) (Fig. 5A). Separation between groups was mainly due to 14 different metabolites (Fig. 5B). Alanine and glycine concentrations remained respectively high and low after 10 days of cyclic hypoxia (Fig. 5C). The effect of cyclic hypoxia on glutamate and BCAAs was no longer detected despite a slightly elevated leucine concentration. The fumarate concentration was high, while the succinate concentration was low in the cyclic hypoxia group.

**Figure 5.**
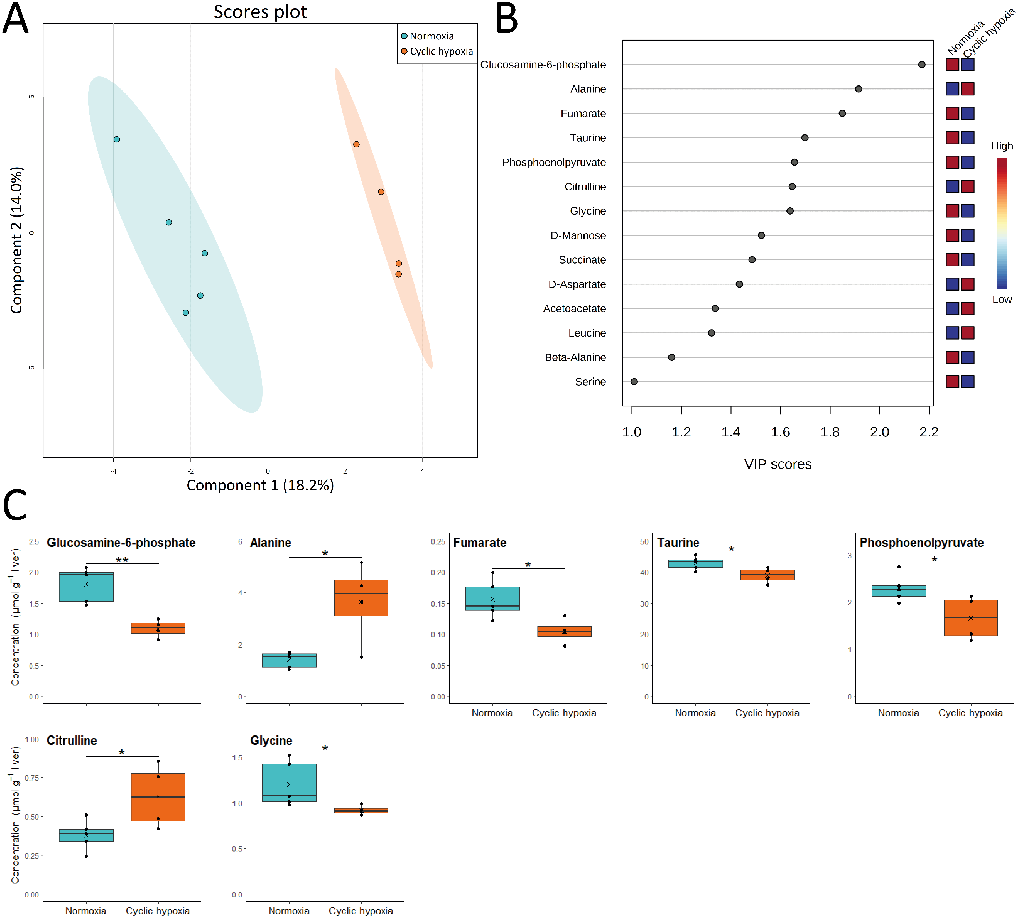
Metabolite profile in liver of Arctic char at day 10 after fish were either maintained in normoxia or exposed to cyclic hypoxia. A) PLS-DA 2D score plots of liver metabolites. Components 1 and 2 represent variance proportion and ellipses correspond to 95% confidence intervals for each treatment group. B) Variable importance in projection (VIP) scores of PLS-DA component 1 for liver metabolites which drive metabolic profile differentiation between the normoxic and cyclic hypoxic groups (VIP score > 1). Relative concentrations of corresponding metabolites are indicated by colored boxes on the right. C) Concentration of metabolites which differ significatively between normoxic and cyclic hypoxic fish. Box plots indicate the median (—), the mean (×), 25^th^ and 75^th^ percentiles (box), 95% range (|) and each observation (●). Two sample t-test results are shown as * (p-value < 0.05), ** (p-value < 0.01).

Finally, after a full month of cyclic hypoxia (D30), the differences between the two groups was down to only 14.1% on the first component (Fig. 6A). Among the 16 metabolites with VIP scores greater than 1.0, two metabolites, succinate and fumarate, especially drove the pattern (Fig. 6B). Both had significantly lower concentrations in fish exposed to cyclic hypoxia (Fig. 6C). Despite the absence of significant differences, the succinate:fumarate ratio had a tendency to decrease after 30 days of cyclic hypoxia (Fig. S3A). Another interesting ratio tendency was pyruvate:lactate (Figure S3B) that showed an upward trend with time in cyclic hypoxia. Conversely, a downward trend was observed in normoxia.

**Figure 6.**
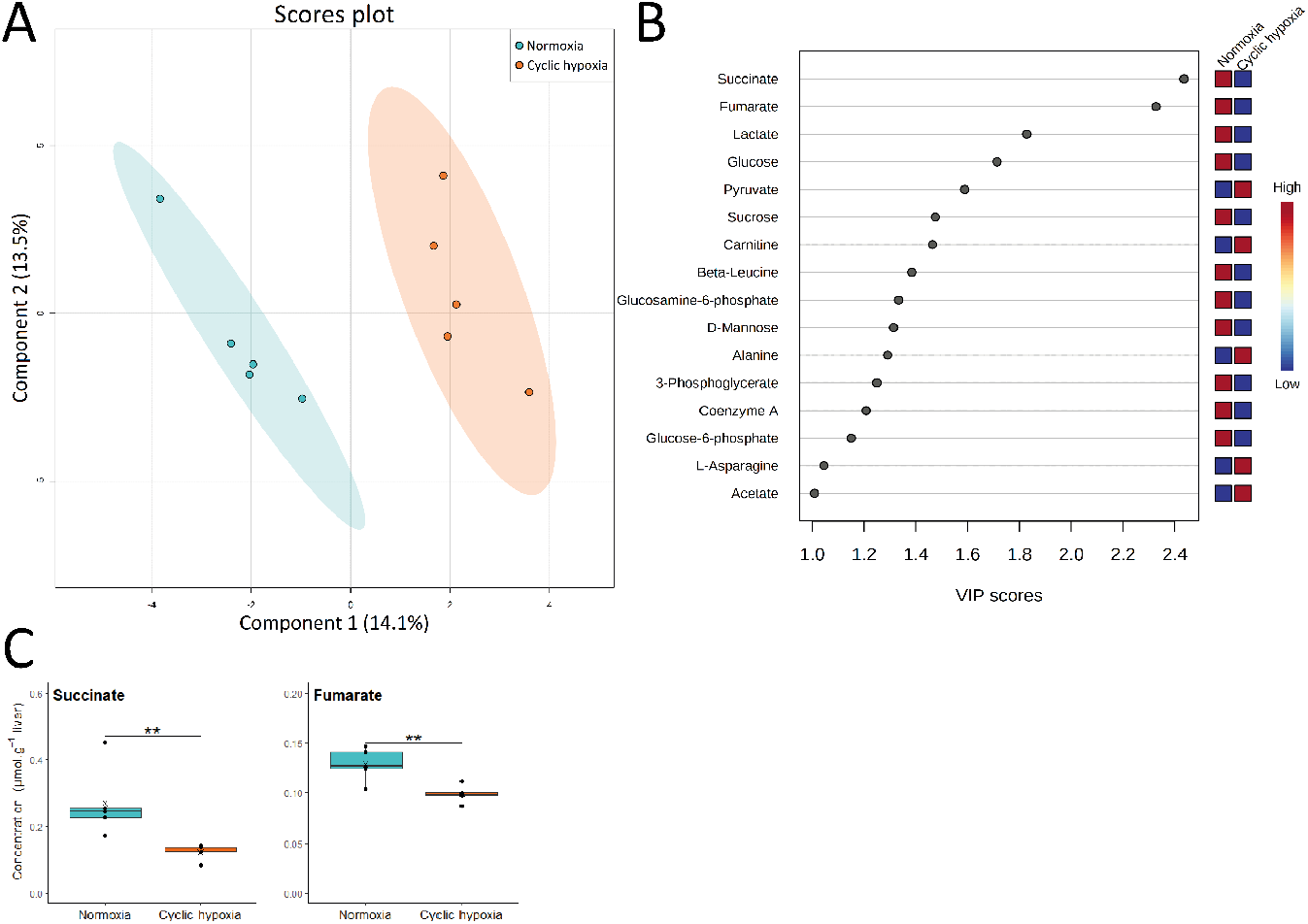
Metabolite profile in liver of Arctic char at day 30 after fish were either maintained in normoxia or exposed to cyclic hypoxia. A) PLS-DA 2D score plots of liver metabolites. Components 1 and 2 represent variance proportion and ellipses correspond to 95% confidence intervals for each treatment group. B) Variable importance in projection (VIP) scores of PLS-DA component 1 for liver metabolites which drive metabolic profile differentiation between the normoxic and cyclic hypoxic groups (VIP score > 1). Relative concentrations of corresponding metabolites are indicated by colored boxes on the right. C) Concentration of metabolites which differ significatively between normoxic and cyclic hypoxic fish. Box plots indicate the median (—), the mean (×), 25^th^ and 75^th^ percentiles (box), 95% range (|) and each observation (●). Two sample t-test results are shown as ** (p-value < 0.01).

## Discussion

This study investigated the effects of cyclic hypoxia on fish growth, energy stores, and metabolism. We hypothesized that cyclic hypoxia would induce a modulation of fish metabolism and a change in energy management that would eventually impair energy allocation toward growth. Our data support a metabolic adjustment in Arctic char liver but revealed a limited impact on growth and energetic stores. Arctic char were able to cope with a month of cyclic hypoxia and maintain their physiological condition through an efficient adjustment of their phenotype.

### Growth and energy stores

Arctic char was more resistant to cyclic hypoxia than anticipated. After one month of cycling, the growth parameters and physiological conditions remained very similar to those of the control group. However, during the first ten days of cyclic hypoxia, the condition of the fish was altered, an observation mainly supported by mass change. This impairment in growth performance due to cyclic hypoxia is also observed in salmon (Remen *et al*., 2014), catfish (Yang *et al*., 2013) and flounder (Davidson *et al*., 2016). In salmon, reduced growth is explained by a decreased food intake but not by food utilization, unlike in catfish. Similar results are observed in seabass and turbot exposed to chronic hypoxia (Pichavant *et al*., 2001). In our experiment, feed ration was adjusted to 0.5% biomass-1 day-1 and no uneaten food was observed. Therefore, it is unlikely that the growth impairment of Arctic char exposed to cyclic hypoxia was caused by reduced food intake. However, it could likely be explained by a decrease in food conversion efficiency or a reduction in energy allocation to growth. This deleterious effect of cyclic hypoxia is compensated for in the following 10 days and completely disappears after about 4 weeks of cyclic hypoxia. Compensatory growth has already been observed in salmon exposed to cyclic hypoxia but only after a one-month recovery period in normoxia (Remen *et al*., 2014). Foss *et al*. reported a recovery from SGR in spotted wolffish exposed to moderate hypoxia for three months (Foss *et al*., 2002). These authors also observed compensatory growth for this species during the recovery period in normoxia (Foss and Imsland, 2002). Compensatory growth reduces risks associated with small size, such as higher mortality, prey limitation, and smaller reproductive success, but comes with associated costs (Ali *et al*., 2003; Mangel and Munch, 2005). In juvenile salmon, compensatory growth is followed by lower long-term performance, which results in smaller size, lower lipid reserve, and reduced incidence of sexual maturation (Morgan and Metcalfe, 2001). A compensatory growth event also induces a decrease of offspring production in guppies (Auer *et al*., 2010) and alters locomotion performance through a reduction of maximum sprint speed in sea bass (Killen *et al*., 2013). However, because of the short period and low intensity of compensatory growth observed in our experiment, it is unlikely that it has long-term negative repercussions. It should be noted that the compensatory growth observed here was not supported by hyperphagia as generally observed (Ali *et al*., 2003).

Arctic char can adjust its metabolism to catch up previously impaired growth. We observed a slightly higher hematocrit under cyclic hypoxia, which may have been coupled with a possible increase in ventilation (although not specifically measured here), thus improving oxygen uptake and delivery to tissues and helping to maintain growth despite hypoxia events. During acclimatation to cyclic hypoxia, we did not detect any noticeable change in tissue energy reserves. White muscle carbohydrates remained unaffected during cyclic hypoxia. In killifish, an increase in skeletal muscle free glucose is observed during the first day of cyclic hypoxia, but not after 28 days of acclimation (Borowiec *et al*., 2018). Consistent with our results, glycogen reserves remain unchanged. The authors reported an increase in muscle lactate up to one hour after the first hypoxic event. In our study, three hours passed between reoxygenation and sampling; this can probably explain the absence of variation in glucose and lactate between the control and experimental groups. In a previous study on the same Arctic char strain, we detected an increase in muscle lactate after 9 hours of exposure to the same level of hypoxia (Cassidy and Lamarre, 2019). After 3 hours of reoxygenation, accumulated muscle lactate should have been excreted into plasma or remobilized into glycogen, as observed in trout (Milligan and Girard, 1993). TAG levels in the liver and visceral fat remained undisturbed throughout the entire experiment. Li *et al*. found that TAG levels decrease in the liver of Nile tilapia, but only after 4 weeks of chronic hypoxia, and that acute hypoxic stress was insufficient to induce the use of TAG in the liver (Li *et al*., 2018). Furthermore, Bera *et al*. reported an increase in plasma TAG levels after 2 months of cyclic hypoxia in goldfish which was interpreted as a suppression of lipolysis or an impairment of TAG incorporation in oocytes (Bera *et al*., 2017). According to our data, TAG and lipid reserves are not used as fuel in Arctic char experiencing cyclic hypoxia.

### Protein synthesis

The rate of hepatic protein synthesis has been shown to curtail during acute hypoxia in Arctic char (Cassidy and Lamarre, 2019). The liver plays a central role in energetic processes, particularly through its involvement in protein metabolism (Charlton, 1996). Therefore, we expected that a compensation of the protein synthesis activity would occur immediately after reoxygenation. We did not observe such a response, suggesting that Arctic char do not accumulate a “protein synthesis debt” during hypoxia. The negligible decrease observed throughout the time in both groups is probably related to the limited feed ration. This would be consistent with previous studies on Arctic char and other species that reported a decreased protein synthesis rate in fasting or starving fish (Lamarre *et al*., 2015; Loughna and Goldspink, 1984; Lowery and Somero, 1990). Although we did not detect significant changes in the rate of protein synthesis, liver metabolism was profoundly affected by cyclic hypoxia, as revealed by metabolomic analysis.

### Metabolite changes in response to cyclic hypoxia

#### Acute stress response

The first hypoxia-reoxygenation cycle caused a profound metabolic reorganisation in the Arctic char liver (Fig. 7A). Amino acid, carbohydrate, and lipid metabolism were particularly impacted. The amino acids accumulated remarkably in the liver of fish exposed to hypoxia. Branched chain amino acids accumulation (leucine, isoleucine, and valine) suggested an increase in protein catabolism combined with a decrease in protein synthesis, as previously described in Arctic char liver during hypoxia (Cassidy and Lamarre, 2019).These amino acids have also been shown to accumulate during anoxia and after up to three hours of reoxygenation in the liver of crucian carp, before decreasing after a full day of recovery (Dahl *et al*., 2021).

**Figure 7.**
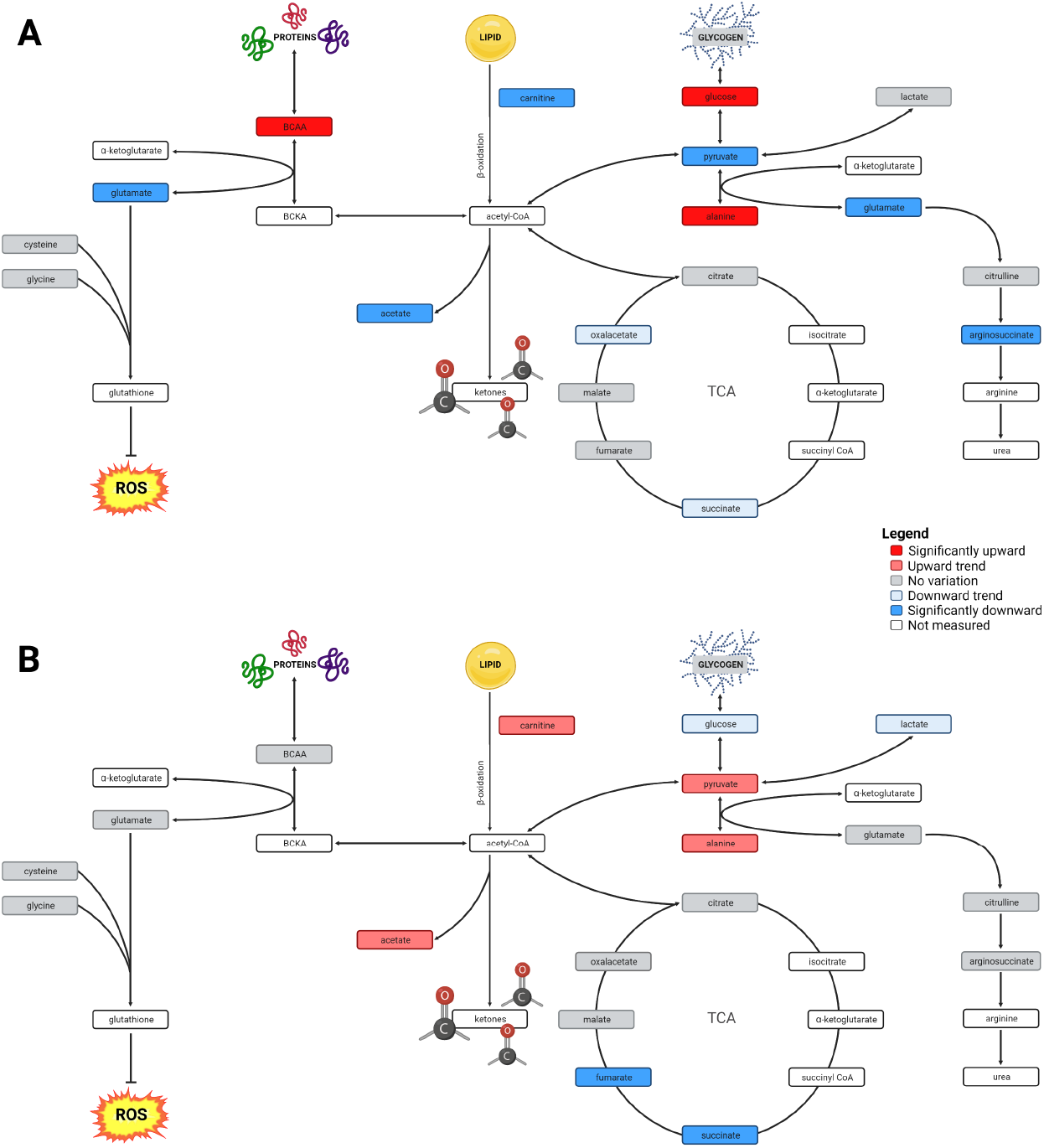
Liver metabolic pathways of Arctic char at day 1 (A) and day 30 (B) after fish were exposed to cyclic hypoxia. Created with BioRender.com.

Alanine is the amino acid that shows the most remarkable changes. It likely accumulated during the hypoxic phase, as demonstrated in the heart and liver of hypoxic frogs (Shekhovtsov *et al*., 2020), anoxic heart of turtles (Bundgaard *et al*., 2019) as well as in the anoxic muscle of common carp, but not in the liver and muscle of hypoxic common carp (Lardon *et al*., 2013). In crucian carp, alanine accumulates in the liver, heart, brain, and plasma during anoxia and its levels remain high up to three hours after reoxygenation (Dahl *et al*., 2021). Wang *et al. (2021)* also measured an increase in alanine aminotransferase activity in yellow catfish liver exposed to six hours of hypoxia, which remained elevated at least six hours after reoxygenation (Wang *et al*., 2021). Our results suggest the activation of the glucose-alanine cycle, also known as the Cahill cycle (Felig, 1973). Alanine, as a product of glycolysis and protein catabolism in muscle, is exported to the liver to be transaminated into pyruvate. This pyruvate is then reduced into glucose, which is exported to muscle to sustain glycolysis. Alternatively, accumulated alanine may also be oxidized and may greatly contribute to liver energy metabolism. Juburi *et al*. (2021) suggested that when available in high concentrations, alanine is the preferred oxidative fuel in trout instead of glucose (Jubouri *et al*., 2021). An alternative hypothesis supported by Bundgaard *et al*. (2019), is that the high NADH/NAD^+^ ratio building up during the lack of oxygen induces a reversal of the TCA cycle. Pyruvate is aminated into alanine to generate α-ketogluratate from glutamate. Oxalacetate is then produced from α-ketoglutarate and aspartate and then reduced into succinate (Bundgaard *et al*., 2019). We did not observe a decrease in aspartate concentration in our study, which could be consistent with another study that observed a recovery in aspartate concentration within three hours of reoxygenation (Dahl *et al*., 2021). Bundgaard *et al*. (2019) also detected stable glutamate concentrations during anoxia, which is consistent with Dahl *et al*. (2021), but not with our results obtained after the reoxygenation. The depletion of glutamate observed here could indicate a regeneration of α-ketoglutarate but not a reversal of the TCA cycle in Arctic char. The most parsimonious explanation for our results is thus that alanine is accumulated in the liver during hypoxia due to muscle exportation and α-ketoglutarate synthesis. During reoxygenation, this alanine is transaminated into pyruvate, which in turn forms glucose via gluconeogenesis, providing an explanation for the concomitant higher hepatic glucose concentration. As mentioned above, glutamate concentration is much lower during acute cyclic hypoxia. Usually, glutamate concentration is maintained or increased in tissues during hypoxia and anoxia (Dahl *et al*., 2021; Shekhovtsov *et al*., 2020) except in the brain, where it generates the neurotransmitter GABA (Dahl *et al*., 2021). Dahl *et al*. (2021) observed a decrease in glutamate concentration in carp liver after a recovery period of one day, but this point was not further discussed. Glutamate is a key metabolite in many metabolic processes. As mentioned, it can supply α-ketoglutarate to the TCA cycle but can also be used to synthesize glutathione, which provides protection against oxidative damages during reoxygenation (Fang *et al*., 2002). However, we did not detect changes in L-cysteine concentration, another amino acid involved in glutathione synthesis. Glutamate is also an important ammonia carrier and can contribute to its excretion in liver through ureagenesis (Li *et al*., 2020). Although it is not the main pathway for ammonia excretion in fish (Wilkie, 2002), glutamate depletion coupled with the decrease in argininosuccinate seem to indicate an eventual elimination of ammonia through urea production. An increase in uric acid and urea in anoxic carp liver previously reported by Dahl *et al*. (2021) may support this hypothesis.

Concerning lipid metabolism, the decline in carnitine levels observed in cyclic hypoxia suggests a decreased β-oxidation capacity, which is also substantiated by decreased acetate levels, a by-product of this pathway (Yamashita *et al*., 2001). Liu *et al*. (2014) demonstrated that hypoxia induced an inhibition of the β-oxidation and promoted lipid accumulation in mouse liver and human hepatocytes (Liu *et al*., 2014). According to Sun *et al. (2020)*, who observed this pattern in largemouth bass, this strategy may help prepare the organism for an eventual long-term hypoxic event (Sun *et al*., 2020).

In general, these observations suggested that the energetic metabolism was under pressure during the first hypoxia-reoxygenation cycle. In addition to the decreased glutamate levels and the accumulation of alanine observed in fish exposed one day to cyclic hypoxia, there was also a depletion of pyruvate concomitant with a slight decrease in oxaloacetate and succinate. The latter is probably accumulated during the hypoxic phase (Chinopoulos, 2019) but is then completely consumed following reoxygenation.

#### Acclimation

After five hypoxia cycles (D5), glutamate, alanine and BCAA levels displayed the same trend as the first day, indicating a relatively similar metabolic response compared to the first day. The slight increase in pyruvate and in the metabolites of the TCA cycle suggest their accumulation during hypoxia but also a lower pressure on energetic metabolism during reoxygenation, in contrast to the acute response. The lower concentration of glycine contrasts with the increase reported in the common carp liver during hypoxia and in the crucian carp liver during anoxia and recovery (Dahl *et al*., 2021; Lardon *et al*., 2013). As a glucogenic amino acid, glycine can contribute to pyruvate production. Glycine is also involved with glutamate in glutathione synthesis and ammonia detoxification in common carp, as shown by Hoseini *et al*. (Hoseini *et al*., 2022). Glycine concentration probably decreased in Arctic char due to the lower capacity of this species to deal with hypoxia compared to common and crucian carps.

After ten days of cycling hypoxia, BCAAs no longer accumulate which may reflect down-regulation of protein degradation during hypoxia. Alanine still accumulated during acclimation to cyclic hypoxia after ten days, but glutamate was not decreased anymore. Additionally, concentrations of metabolites of the TCA cycle that were slightly elevated at D5 returned to control concentrations after 10 days of cyclic hypoxia. Interestingly, fumarate and succinate concentrations are lower after ten days of cyclic hypoxia. When succinate accumulated during hypoxia is oxidized to fumarate during reoxygenation, reactive oxygen species (ROS) are generated (Chouchani *et al*., 2014). Keeping the concentration of succinate in check during hypoxia may be a strategy to reduce oxidative stress during cyclic hypoxia. This mechanism is even more obvious after one month of cyclic hypoxia (D30). Both succinate and fumarate concentrations are lower in the hypoxic group and are the only metabolites that differ in the two groups (Fig. 7B). Furthermore, we observed a reduced, albeit not significant, succinate:fumarate ratio in fish exposed to cyclic hypoxia (Fig. S3B). Keeping this ratio low may be viewed as a protective mechanism against ROS. Hypoxia-tolerant species tend to have a lower succinate:fumarate ratio during hypoxia exposure (Bundgaard *et al*., 2019; Shekhovtsov *et al*., 2020).

Finally, a slightly higher pyruvate:lactate ratio observed in the cycling fish indicates a decreased dependence on anaerobic glycolysis. In general, after 30 days of cycling hypoxia, both groups of fish showed a similar metabolome, suggesting that Arctic char exposed to one month of cyclic hypoxia successfully adopted a phenotype that is more tolerant to cyclic hypoxia.

To conclude, this study reveals that Arctic char can acclimate well to cyclic hypoxia. Despite the small growth impairment observed in the first days, fish quickly recovered and maintained the gain in mass and length while defending their energy stores. Metabolic disturbances occurred mostly after the first few hypoxia events. The liver metabolome is profoundly altered at the level of carbohydrate and amino acid metabolism during acute hypoxia cycles but after one month of cyclic hypoxia, most of these effects are resolved, supporting the adoption of a new phenotype.

As reported by Anttila *et al*., Arctic char demonstrated great hypoxia resistance for a salmonid (Anttila *et al*., 2015). Our study confirms the resilience of this species when faced with oxygen level variations. More studies, specifically in transcriptomics or proteomics, would be necessary to understand the nature of this new phenotype. We point out that the first cyclic hypoxia cycles are particularly demanding for Arctic char, which need several days to adjust their physiology. This period could thus be critical if fish face other stress events at the same time, such as high temperature or predation. In this context, Arctic char populations living in shallow waters may be challenged during summer with the present increase of extreme climate events.

## List of symbols and abbreviations

CF: condition factor
DSS: 4,4-dimethyl-4-silapentane-1-sulfonic acid
ETS: electron transport system
FP: protein-free amino acid pool
*Ks*: fractional rate of protein synthesis
PCA: perchloric acid
PFBBr: pentafluorobenzyl bromide
PHE: phenylalanine
PLS-DA: partial least squares discriminant analysis
PP: protein pool
ROS: reactive oxygen species
SGR: specific growth rate
TAG: triglycerides
VIP: variable importance to the projection

## Acknowledgements

We thank Emilie Bourloutski, Antoine Zboralski, Chloé Melanson, Claude Power, and Adrien Biessy for their help with sampling sessions and Robert Cormier for his support in using the Chenomx software.

